## Appendix (Supplemental) Figures for "Mechanism and Structure-Guided Optimization of SLC1A1/EAAT3-Selective Inhibitors in Kidney Cancer"

**for**

**Table of Contents	Page No.**

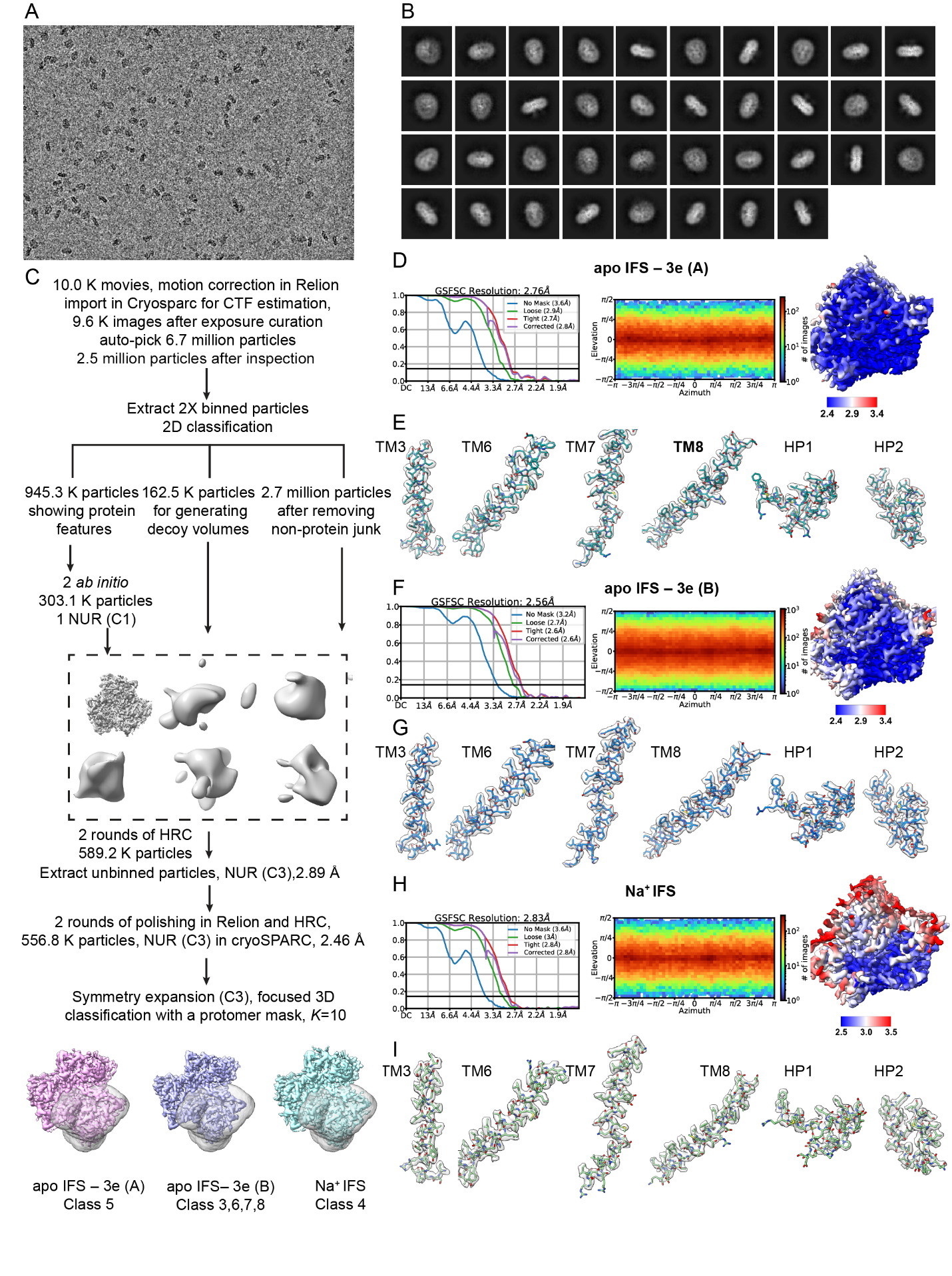

**Appendix Figure S1. Cryo-EM data processing and refinement.** A representative image (**A**) and selected 2D class averages (**B**) of the dataset. (**C**) The cryo-EM data processing flow, showing identification of three protomer structural classes – two with bound 3e (classes A and B) and one bound to Na^+^ ions. (**D-I**) The golden standard Fourier shell correlation (FSC) curves of the final refinement (**D, F, H, left**), the angular distribution of particles used for the final 3D reconstitutions (**D, F, H, middle**), the local resolution distribution (**D, F, H, right**), and the EM density of transport domain transmembrane helices (TMs), and helical hairpins (HPs) (**E, G, I**) for structural class A of *apo* IF state with **3e** (**D, E**), class B of *apo* IF state with **3e** (**F, G**), and Na^+^-bound IF state (**H, I**). The map contour levels of (**E**), (**G**), and (**I**) in ChimeraX are 0.71, 0.732, and 0.654, respectively, corresponding to 5σ.

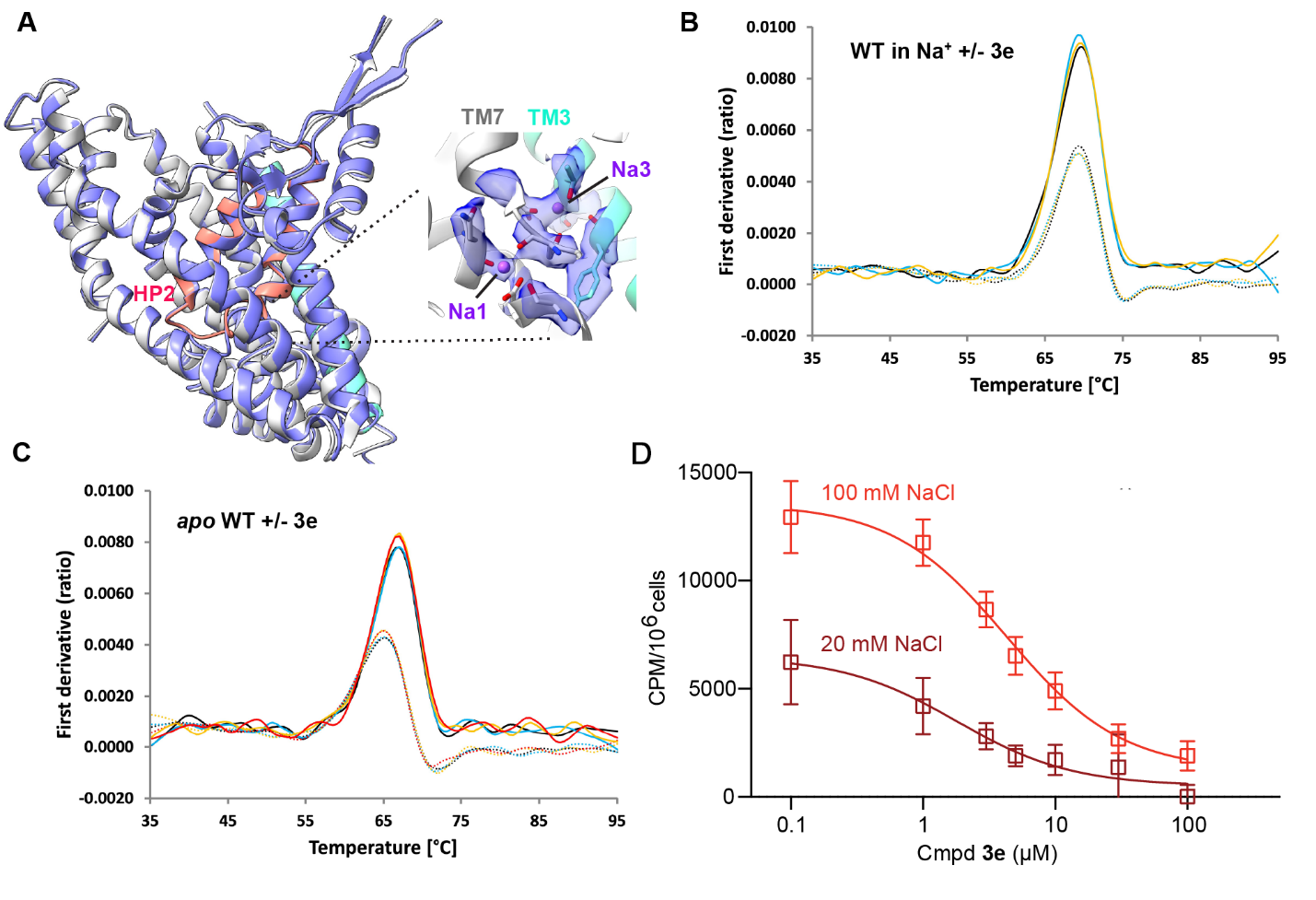

**Appendix Figure S2. NanoDSF measuring WT SLC1A1 and compound 3e binding with and without Na^+^.** (**A**) The Na^+^-bound monomer observed in the presence of **3e** and refined to 2.83 Å (gray with HP2 in pink and TM3 in cyan) is indistinguishable from Na^+^-bound SLC1A1g (state blue, PDB accession code 6x2l) imaged in the absence of **3e**; the entire protein was superposed. The inset shows the enlarged Na1 and Na3 sites with well-resolved EM density (contoured at 5 σ) for the bound Na^+^ ions. Coordinating residues are shown as sticks. (**B, C**) NanoDSF traces showing the first derivative of the fluorescence emission ratio at 350 and 330 nm of the WT SLC1A1g in 200 mM Na^+^ (**B**) or under *apo* conditions in buffer containing 100 mM NMDG (**C**) without (dashed lines) and with (solid lines) 100 µM **3e.** (**D**) L-[^3^H]-Asp uptake into cells expressing SLC1A1g in the presence of 100 (red) and 20 mM (brown) NaCl. Raw counts per 10^6^ cells are shown; background counts in unresected cells averaged at ~1700±100 CPM were subtracted from the data. The fitted IC_50_ values are 4.3 ± 0.5 and 1.8 ± 0.7 µM, respectively.

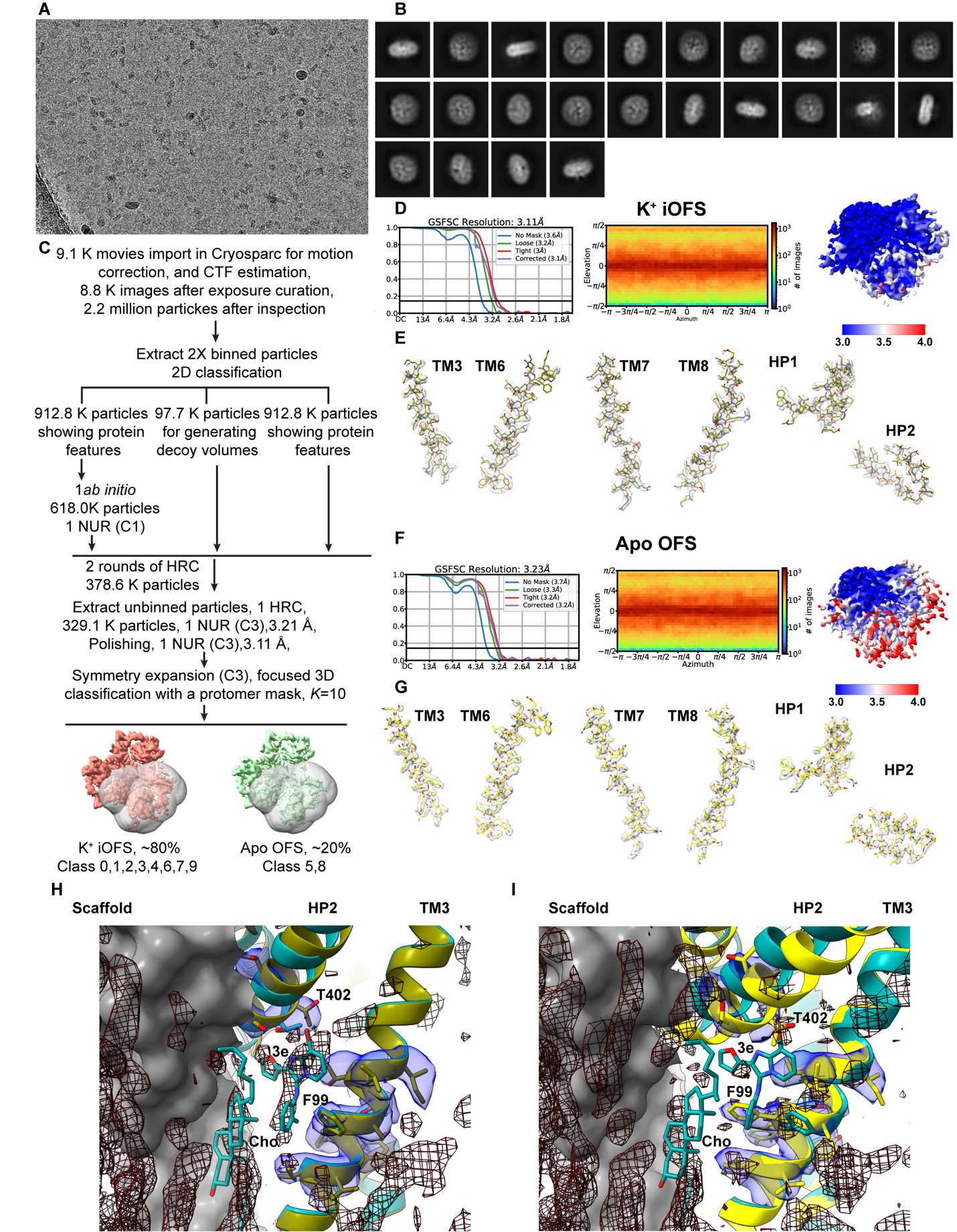

**Appendix Figure S3. Cryo-EM structure of Hg2+-crosslinked K269C/W441C SLC1A1g mutant in the presence of 100 μM 3e.** A representative image (**A**) and selected 2D class averages (**B**). (**C**) The cryo-EM data processing flow, showing the identification of two protomer structural classes: an intermediate bound to a K^+^ ion (K^+^ iOFS) and the *apo* OF state (Apo OFS). (**D-G**) The golden standard Fourier shell correlation (FSC) curves of the final refinement (**D, F, left**), the angular distribution of particles used for the final 3D reconstitutions (**D, F, middle**), the local resolution distribution (**D, F, right**), and the EM density of transport domain transmembrane helices (TMs), and helical hairpins (HPs) (**E, G**) for K^+^ iOFS state and Apo OFS. The map contour levels in ChimeraX are 0.098 (**E**) and 0.076 (**G**), corresponding to 4σ. (**H, I**) Molecular models (PDB IDs 8CUA, olive; and 8CUD, yellow) were rigid-body fitted without further adjustment into the density of K iOFS (3.11 Å resolution; **H**) and the Apo OFS (3.22 Å resolution; **I**. The transport domain from the **3e**-bound structure (teal) is superimposed for comparison using residues 84-124 and 275-475. TM6, HP1, and TM7 are omitted for clarity; residues F99, T402, compound **3e**, and cholesterol (Cho) are shown as sticks. The scaffold domain is displayed as a gray surface. Protein density in the vicinity of F99 and T402 is shown as a blue surface, whereas non-protein density is shown as a black mesh. No well-defined density corresponding to **3e** or cholesterol is observed, although a weak, elongated density is present at the **3e**-binding site in the K^+^-bound intermediate state (**H**). Conformational shifts in HP2 disrupt interactions between T402 and **3e** and introduce steric clashes in the K^+^-bound intermediate and *apo* OF transport domains, respectively. Map contour levels in ChimeraX are 1.23 (**H**) and 0.95 (**I**), corresponding to 5σ.

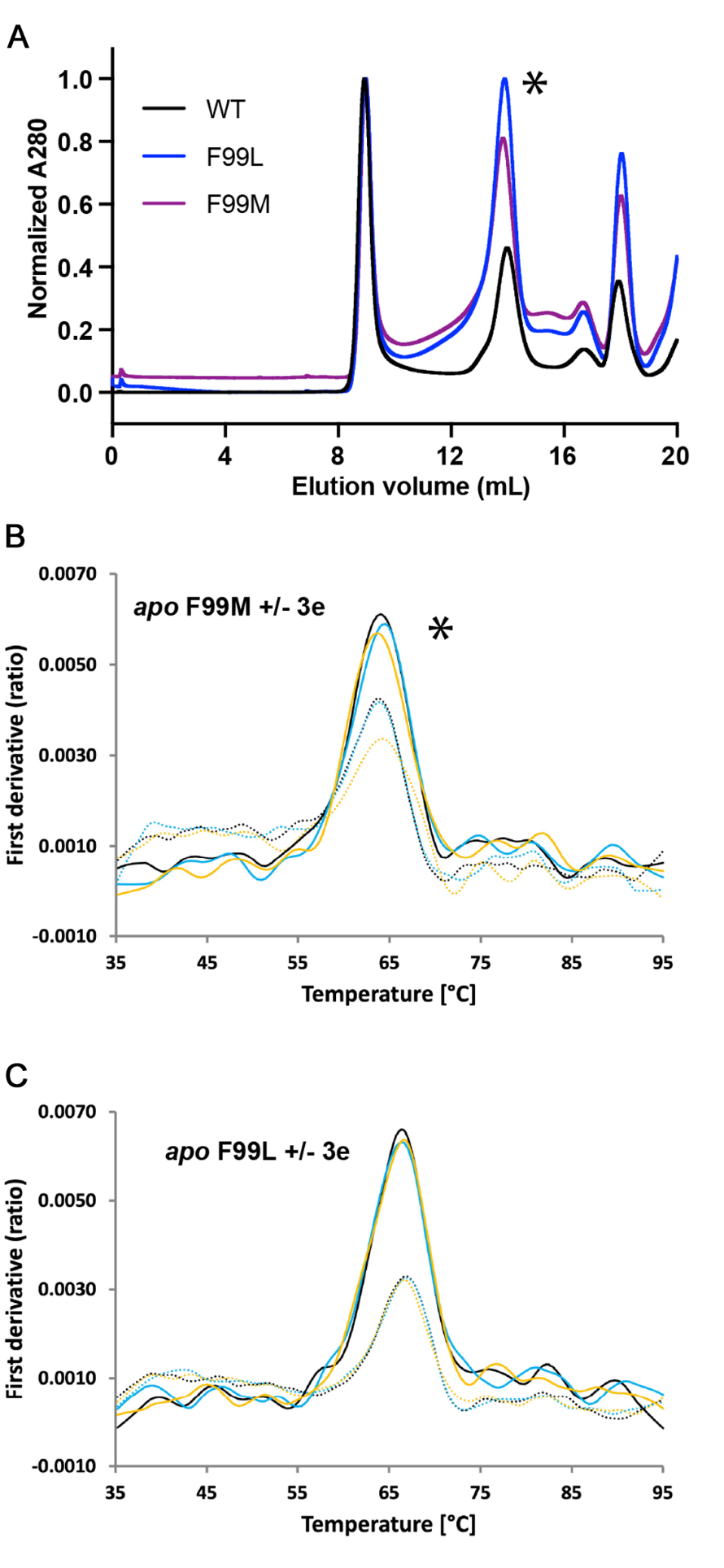

**Appendix Figure S4. NanoDSF Measuring WT and mutant SLC1A1 purification and thermal stability.** (**A**) Preparative size exclusion chromatography of the WT SLC1A1g and drug-resistant mutants F99L and F99M. An asterisk indicates the elution peak containing the trimeric protein, which was collected and used for nanoDSF experiments. (**B, C**) NanoDSF traces for the drug-resistant F99M (**B**) and F99L (**C**) mutants were recorded under *apo* conditions without (dashed lines) and with (solid lines) 100 µM **3e**. An asterisk indicates an experiment where two biological repeats exhibited slightly different absolute *Ti* values, while yielding similar *ΔTi*.

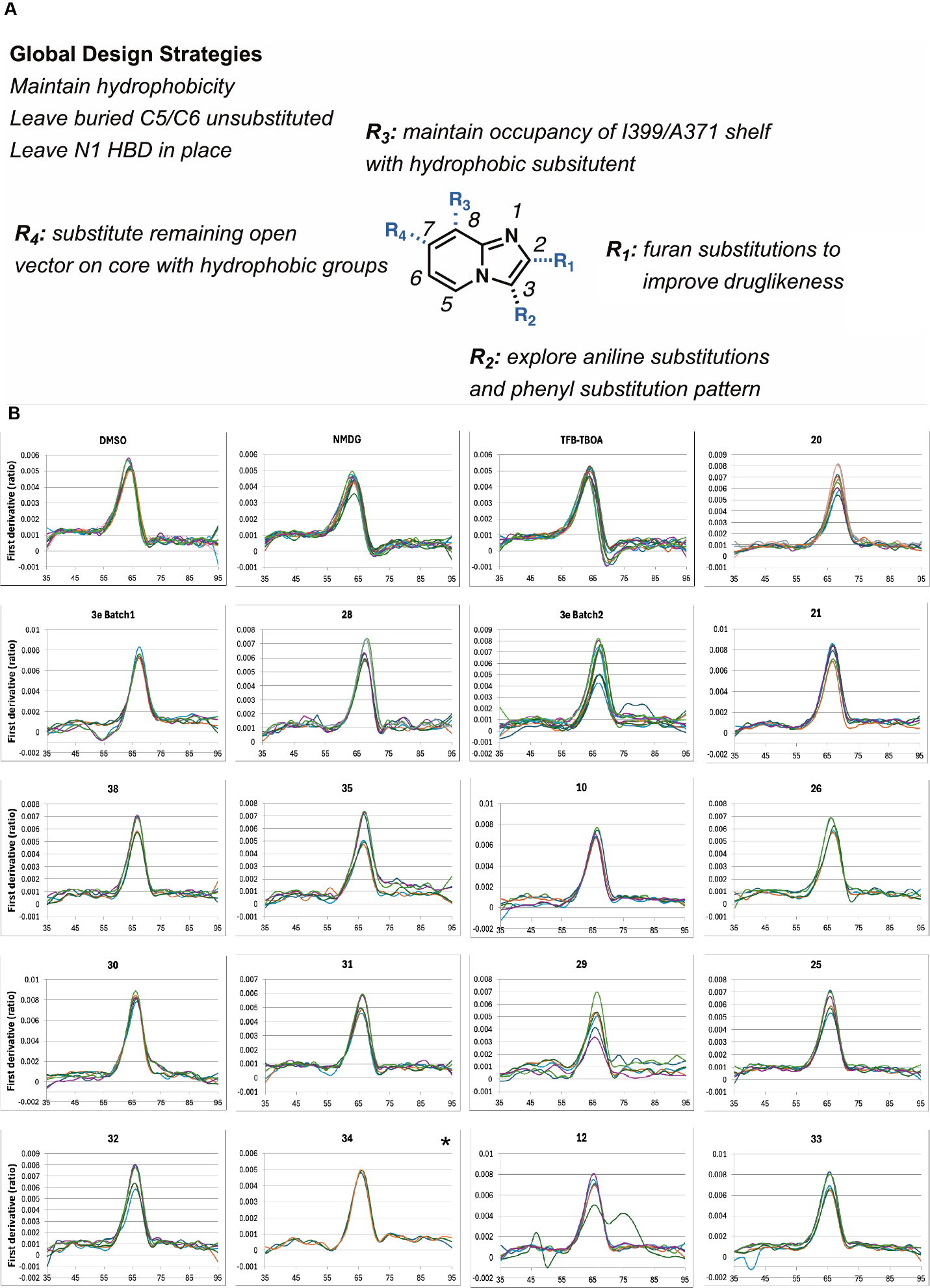

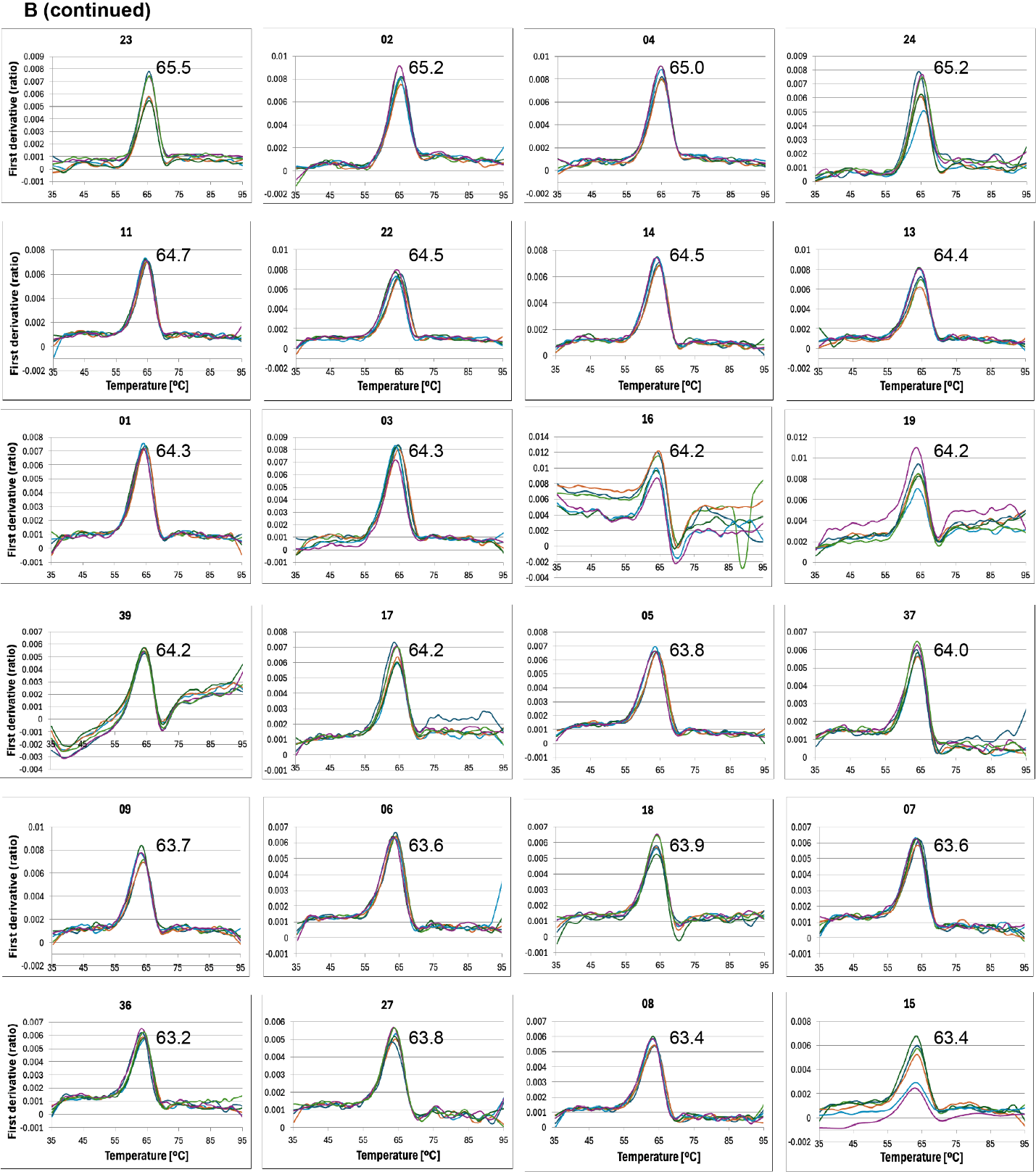

**Appendix Figure S5. BIA analog design and raw nanoDSF data for BIA compound screening.** (**A**) Optimization strategy of BIA analogs based on compound **3e**. (**B**) NanoDSF traces showing the first derivative of the fluorescence emission ratio at 350 and 330 nm of the SLC1A1g. The data were recorded under *apo* conditions in the presence of 200 mM NMDG chloride (NMDG) with additions of 10% DMSO or TFB-TBOA, which does not bind without Na^+^ ions, as controls, and 100 µM of **3e** (two independently prepared batches) and its analogs, as indicated above the graphs. The inflection temperature (*T_i_)* was determined from the maximum value of the peaks. The average *T_i_*-s are shown on all plots. All measurements were performed using two independent protein preparations, each with at least three technical replicates. Each line in the graph represents an individual replicate. An asterisk (*) indicates a condition in which one biological replicate yielded an abnormal unfolding profile, excluded from the analysis.

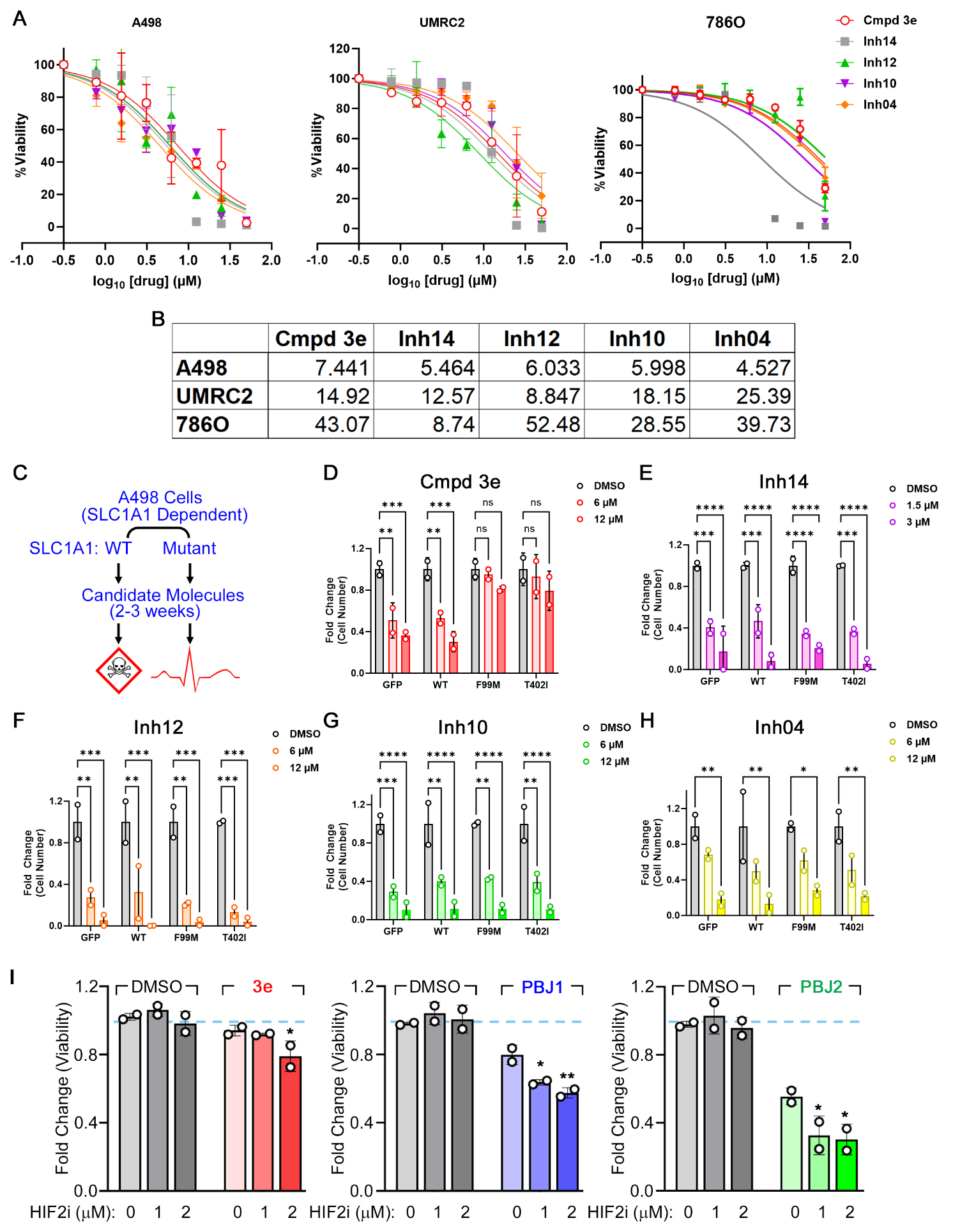

**Appendix Figure S6. On-target validation of candidate BIA analogs in cell-based assays.** (**A** and **B**) Cell viability measurement of the indicated RCC cells (**A**) and a table with the IC_50_ values, calculated using regression analysis in Graphpad Prism (**B**), upon treatment with the indicated compounds for 7 days. (**C**) Schema depicting the experimental design to establish on-target effects of cmpd **3e** analogs using the SLC1A1 mutants. (**D** to **H**) Fold change in cell number, plotted relative to cell counts in the DMSO-treated arm, of A498 cells expressing the indicated SLC1A1 constructs or GFP control, treated with the indicated concentrations of **3e** (**D**), **14** (**E**), **12** **(F**), **10** (**G**), and **04** (**H**), for 21 days. Cell counts, measured using ViCell, were compared using ANOVA (relative to GFP within each group) with Dunnett’s multiple comparison test, n=2, *p<0.05, **p<0.01, ***p<0.001, ****p<0.0001, ns=non-significant. (**I**) Fold change in cell counts, calculated after 7 days of exposure, in A498 cells that were treated with either 12 μM of the indicated SLC1A1 inhibitor or DMSO (control), in the presence or absence of the indicated concentrations of the HIF2α inhibitor PT2385.

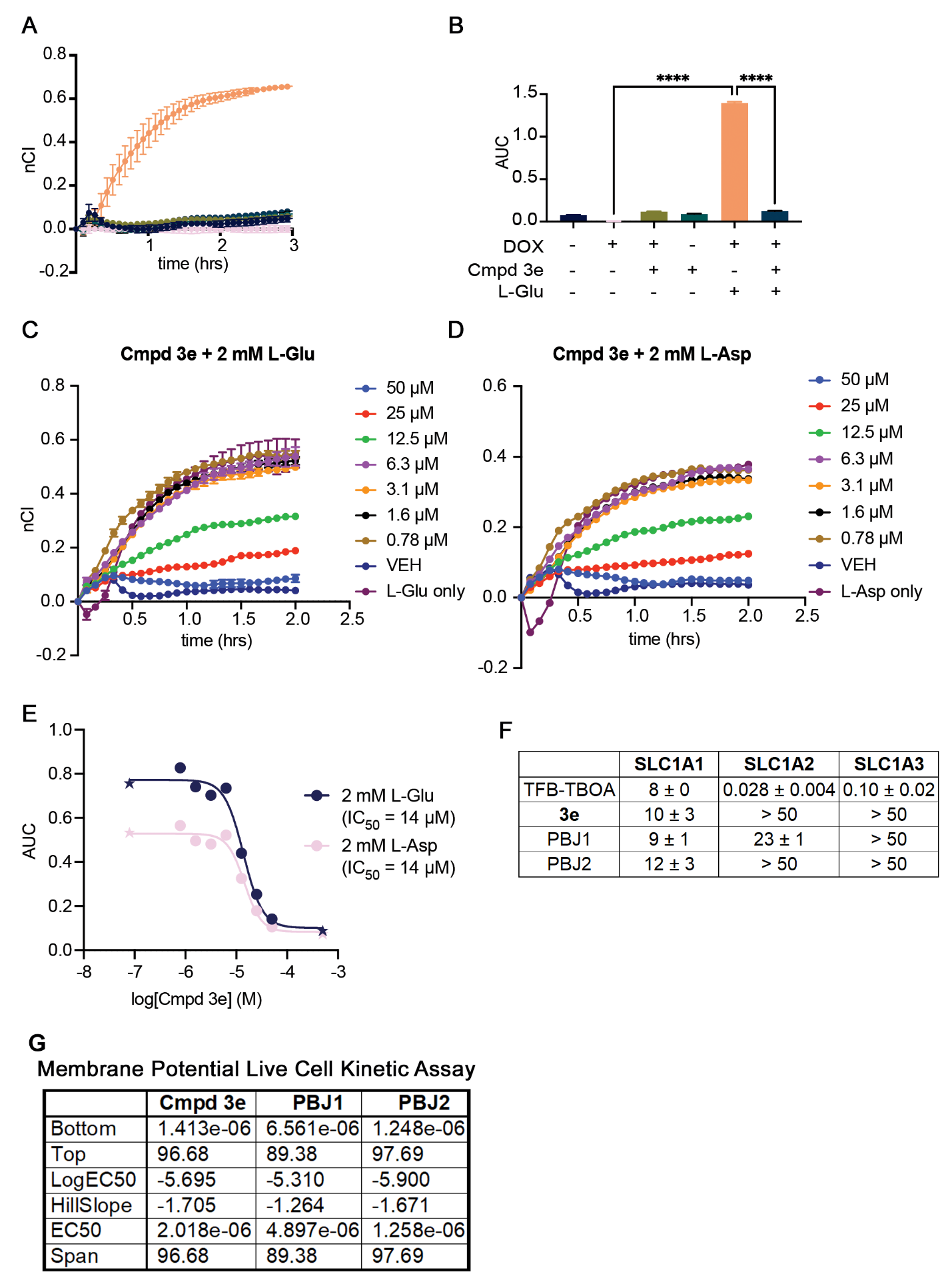

**Appendix Figure S7. Compound 3e inhibits SLC1A1-mediated uptake of both L-Glu and L-Asp. (A**-**B**) Normalized cell index (nCI) (**A**) and integrated Are Under the Curve (AUC) (**B**) for HEK293-SLC1A1 cells treated with 0.15 µg/mL doxycycline (DOX), 1 mM L-Glu, and/or 50 µM **3e** during an xCELLigence impedance assay. HEK293-SLC1A1 cells respond to L-Glu with an increase in nCI when SLC1A1 expression is induced with Dox (orange), but the cells do not respond to L-Glu when Dox is omitted (green). The addition of **3e** inhibits the Dox-dependent morphological response to L-Glu (navy). (**C**-**E**) Impedance based measurement of nCI (**C** and **D**) or integrated AUC (**E**) in HEK293-SLC1A1 cells that were treated with the indicated doses of cmpd **3e**, or vehicle (VEH) control, either in the presence of 2 mM L-Glu or 2 mM L-Asp, as indicated. Data are presented as mean ± S.D. from two technical replicates. (**F**) IC_50_ values measured from the impedance assays, described in main figure 6D. (**G**) IC_50_ values measured from the membrane potential FLIPR assays, described in main figures 6E and 6F.

**Appendix Table S1. Cryo-EM data collection, refinement, and validation statistics**

|  | *apo* IFS - **3e**  (A) | *apo* IFS - **3e**  (B) | Na^+^ IFS | K^+^ iOFS | Apo OFS |
| --- | --- | --- | --- | --- | --- |
| **Data collection and processing** |  |  |  |  |  |
| Magnification | 100,500 X | | | 81,000 X | |
| Voltage (kV) | 300 | | | 300 | |
| Electron exposure (e–/Å^2^) | 58 | | | 53.9 | |
| Defocus range (μm) | -1.0 – -2.0 | | | -0.8 – -2.5 | |
| Pixel size (Å) | 0.83 | | | 0.856 | |
| Initial particle images (no.) | 2,717,842 (1,470,412, C3 expanded particles) | | | 2,187,170 (987,213) | |
| Symmetry imposed | C1 | C1 | C1 | C1 | C1 |
| Final particle images (no.) | 165,757 | 659,564 | 165,684 | 791,674 | 195,539 |
| Map resolution (Å)  FSC threshold | 2.76  0.143 | 2.56  0.143 | 2.83  0.143 | 3.11  0.143 | 3.23  0.143 |
| Map resolution range (Å) | 39.45 – 2.43 | 32.54 – 1.81 | 40.85 – 2.47 | 37.15 – 1.83 | 40.87 – 2.64 |
| **Refinement** |  |  |  |  |  |
| Initial model used (PDB code) | 6X3F | 6X3F | 6X2L | This map is similar to EMD-26997. The model 8CUA can be fitted in this map. | This map is similar to EMD-26998. The model 8CUD can be fitted in this map. |
| Model resolution (Å)  FSC threshold | 2.9  0.5 | 2.7  0.5 | 3.1  0.5 |  |  |
| Map sharpening *B* factor (Å^2^) |  |  |  |  |  |
| Model composition  Non-hydrogen atoms  Protein residues  Ligands | 3,224  412  3 | 3,238  409  4 | 3,167  407  4 |  |  |
| *B* factors (Å^2^)  Protein  Ligand | 46.66  52.58 | 35.81  40.91 | 47.25  48.34 |  |  |
| R.m.s. deviations  Bond lengths (Å)  Bond angles (°) | 0.005  0.949 | 0.005  1.007 | 0.004  0.928 |  |  |
| Validation  MolProbity score  Clashscore  Poor rotamers (%) | 1.23  4.51  0.00 | 1.02  2.39  0.00 | 1.21  4.27  0.00 |  |  |
| Ramachandran plot  Favored (%)  Allowed (%)  Disallowed (%)  **PDB code**  **EMDB code** | 98.03  1.97  0.00  9P4X  71288 | 99.01  0.99  0.00  9P4Y  71289 | 98.25  1.75  0.00  9P4Z  71290 | 75048 | 75049 |

**Appendix Table S2. Structure activity relationship (a)**

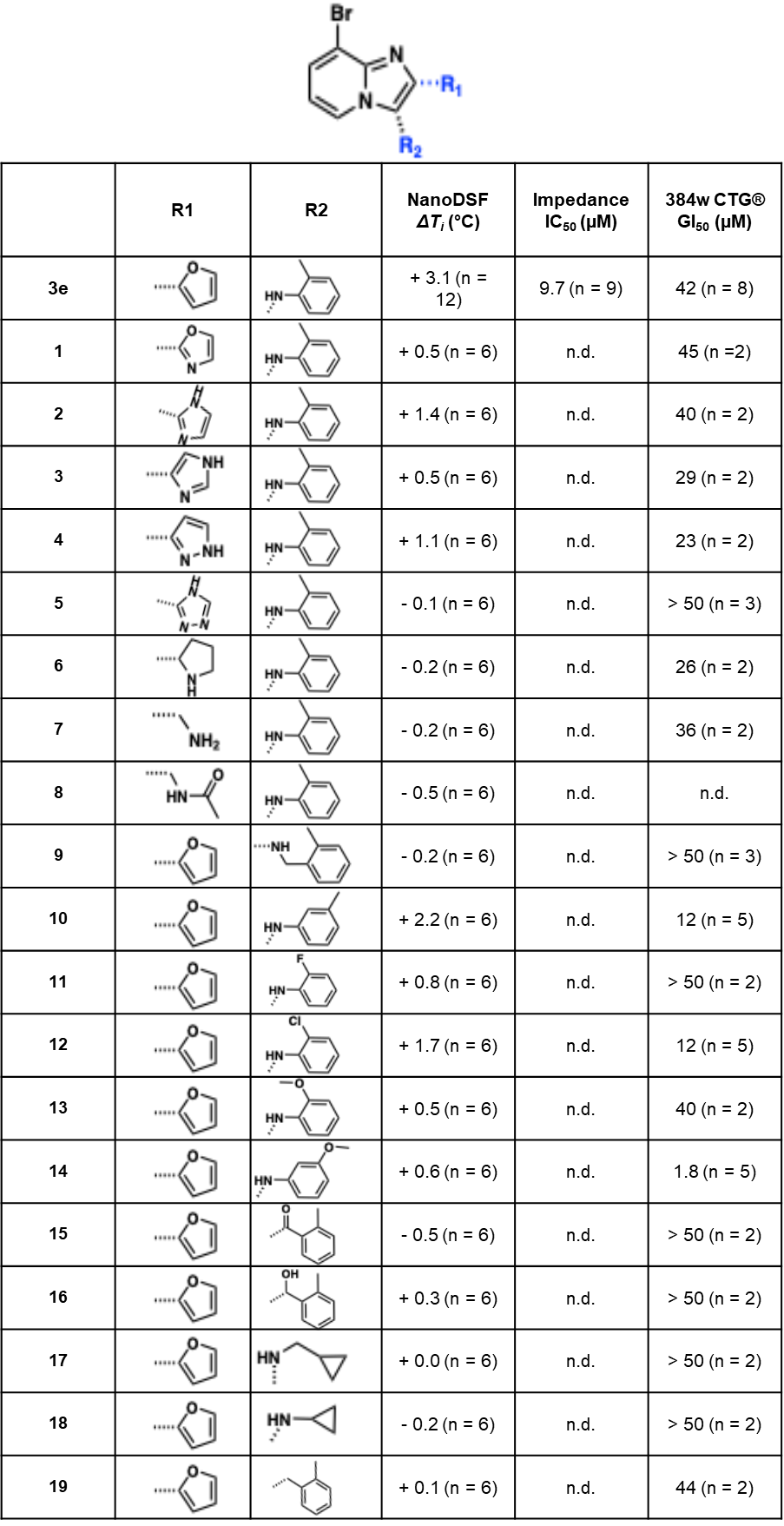
**Appendix Table S2. Summary data of BIA analog SAR.** Analog compound numbers with R_1_/R_2_ modifications on a constant 8-bromo core, chemical structures, NanoDSF thermal shifts (*ΔT_i_*) reflecting binding to SLC1A1g, IC_50_ values calculated using the HEK293-SLC1A1 impedance assays, and GI_50_ values measured using Cell-Titer Glo. Data are presented as mean ± S.D. from the indicated number of biological replicates (n).

**Appendix Table S3. Structure activity relationship (b)**
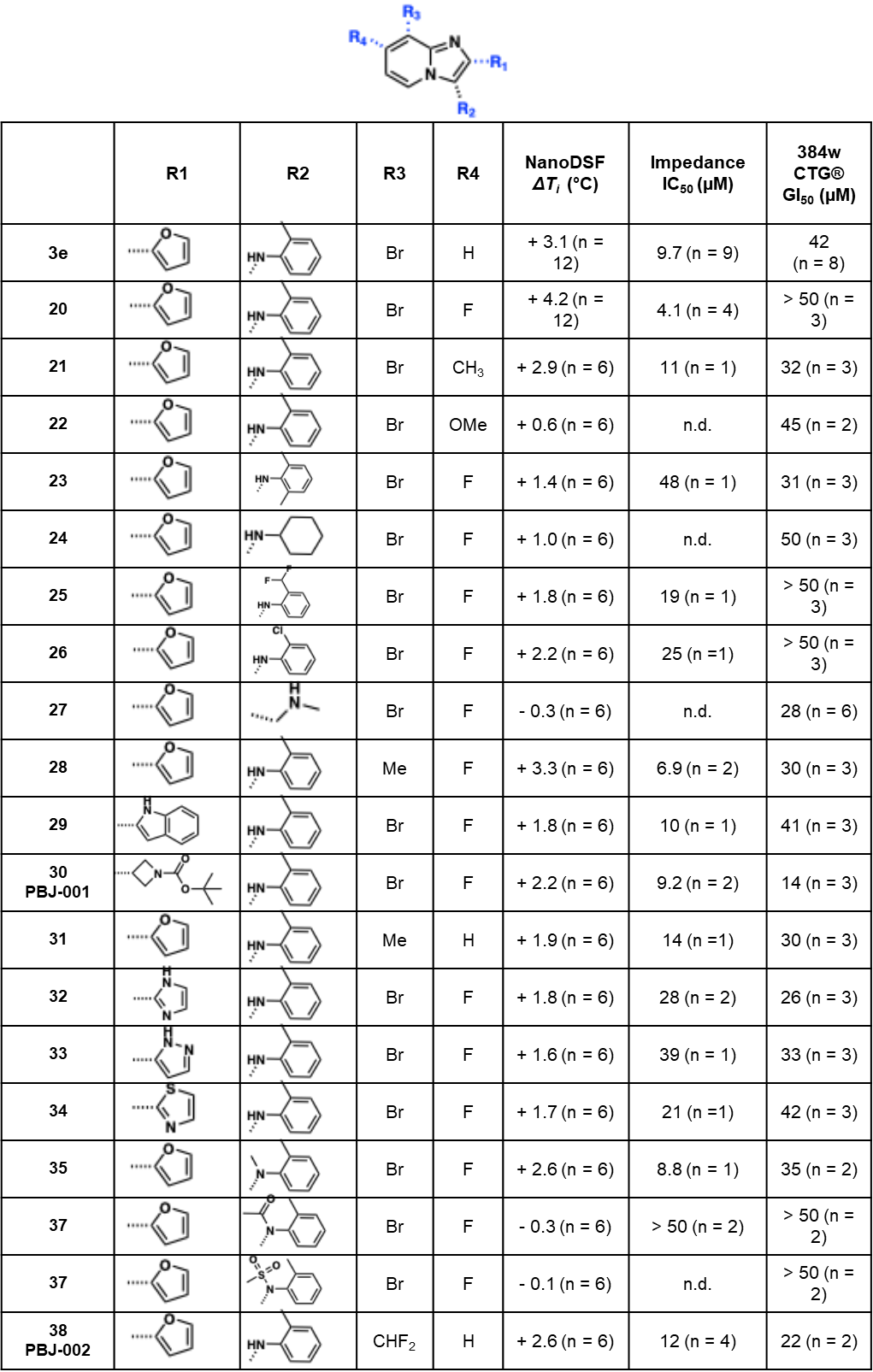

**Appendix Table S3. Summary data of BIA analog SAR.** Analog compound numbers and chemical structures of **3e** analogs with the indicated modifications in R_1_—R_4_, NanoDSF thermal shifts (*ΔT_i_*) reflecting binding to SLC1A1g, IC_50_ values calculated using the HEK293-SLC1A1 impedance assays, and GI_50_ values measured using Cell-Titer Glo. Data are presented as mean ± S.D. from the indicated number of biological replicates (n).
